# Causal gene regulatory analysis with RNA velocity reveals an interplay between slow and fast transcription factors

**DOI:** 10.1101/2022.10.18.512766

**Authors:** Rohit Singh, Alexander P. Wu, Anish Mudide, Bonnie Berger

## Abstract

Single-cell expression dynamics from differentiation trajectories or RNA velocity have the potential to reveal causal links between transcription factors (TFs) and their target genes in gene regulatory networks (GRNs). However, existing methods either neglect these expression dynamics or require cells to be ordered along a linear pseudotemporal axis, which is incompatible with branching trajectories. We introduce Velorama, an approach to causal GRN inference that represents single-cell differentiation dynamics as a directed acyclic graph (DAG) of cells constructed from pseudotime or RNA velocity measurements. In contrast to previous approaches, Velorama is able to work directly with RNA velocity-based cell-to-cell transition probabilities and enables estimates of TF interaction speeds with their target genes. On a set of synthetic datasets, Velorama substantially outperforms existing approaches, improving area under the precision-recall curve (AUPRC) by 3.7–4.8x over the next best method. Applying Velorama to four RNA velocity datasets, we uncover evidence that the speed of a TF’s interactions is tied to its regulatory function. For human corticogenesis, we find slow TFs to be linked to gliomas and co-regulate preferentially with fast TFs, while fast TFs are associated with neuropsychiatric diseases. We expect Velorama to be a critical part of the RNA velocity toolkit for investigating the causal drivers of differentiation and disease.

**Software availability:** https://cb.csail.mit.edu/cb/velorama

## 1 Progress and Potential

A long-standing goal in molecular biology has been to map how the thousands of genes in the human genome interact with each other to determine cellular function. These maps, often termed gene regulatory networks, can provide essential information about which genes drive key stages of cellular differentiation and are major determinants of disease. However, initial efforts to construct gene regulatory networks were limited by technologies that could only account for entire populations of cells altogether but not individually. These bulk readouts often obscured critical but nuanced patterns of gene expression that characterize how cells transition from one state to the next. Recent advances, though, have enabled the measurements of gene expression for individual cells. In addition to capturing more subtle gene expression patterns, these measurements also capture cellular dynamics by treating each single-cell observation as a snapshot of dynamic cell state. Just as a series of photos taken over time can depict movement, these single-cell snapshots can be stitched together to capture differentiation dynamics, oftentimes revealing complex trajectories with multiple branches. However, existing approaches often struggle to handle such complex cell-state landscapes since their mathematical foundations require a sequential ordering of cells. A crucial challenge, therefore, is to determine how to analyze single-cell data in a manner that not only portrays the multitude of possible cell behaviors but also exploits these dynamics to predict cause-effect relationships between genes.

The key conceptual advance of Velorama is to leverage a flexible single-cell graph representation to infer causal gene regulatory networks that can capture complex dynamics. Velorama takes a temporal causality approach, searching for lagged interactions between transcription factors and their target genes. We focus on RNA velocity, a powerful concept that uses splicing dynamics to predict the forthcoming state each cell is likely to adopt. While previous approaches have struggled to interpret the complex dynamics encoded in RNA velocity readouts, Velorama directly incorporates per-cell RNA velocity estimates into a graph of cells, flexibly representing complex trajectories. Velorama’s neural network formulation infers causality using this graph, substantially outperforming state-of-the-art approaches for gene regulatory network inference. Importantly, by explicitly focusing on causal interactions that feature lagged interactions, Velorama also newly enables inquiries into the role of timing and speed in gene regulatory mechanisms. When applied to human neural development, it uncovered important links between regulatory speed and disease, associating gliomas and neuropsychiatric diseases with slow and fast regulators, respectively. These findings point to the potential of Velorama to unlock fundamental insights into cellular differentiation and disease that were previously obscured. With the ever growing quantities of single-cell datasets and ongoing progress in modeling cellular dynamics, Velorama is well-positioned to serve a vital role in advancing our understanding of a wide range of gene regulatory processes.

## 2 Introduction

Understanding the mechanisms that drive cellular differentiation and cell state transitions requires the inference of gene regulatory networks (GRNs) that capture causal transcription factor (TF)–target gene interactions [1–3]. Single-cell transcriptomic technologies, like single-cell RNA-seq (scRNA-seq), have unlocked enormous potential for inferring GRNs that represent these dynamic biological processes. By providing transcriptomic profiles for individual cells, scRNA-seq enables the discovery of cell states that previously would have been obscured with bulk sequencing [4]. We focus here on a powerful innovation in scRNA-seq-based GRN inference that takes advantage of scRNA-seq datasets’ ability to capture snapshots of cell states along dynamic biological processes. These snapshots of cell states can be computationally ordered to infer “pseudotime” stamps for cells that approximate their relative position along a process. Recent GRN inference methods exploit this pseudotime-based ordering of cells to compare the expression profile of a gene against the lagged profile of its putative TF regulators [5–11]. This ordering of cells enables more accurate inference and brings us closer to discovering causal regulatory relationships rather than just gene associations.

Unfortunately, existing GRN inference methods that incorporate pseudotemporal information have a fundamental limitation. They require a *total*, or linear, ordering over cells (or groups of cells) in order to create a temporal axis along which each gene’s expression profile can be correlated with expression profiles of other genes. For a trajectory with multiple branches, a total ordering of cells would collapse these distinct branches down into a single global sequence of cells (**Figure 1A**).

**Figure 1:**
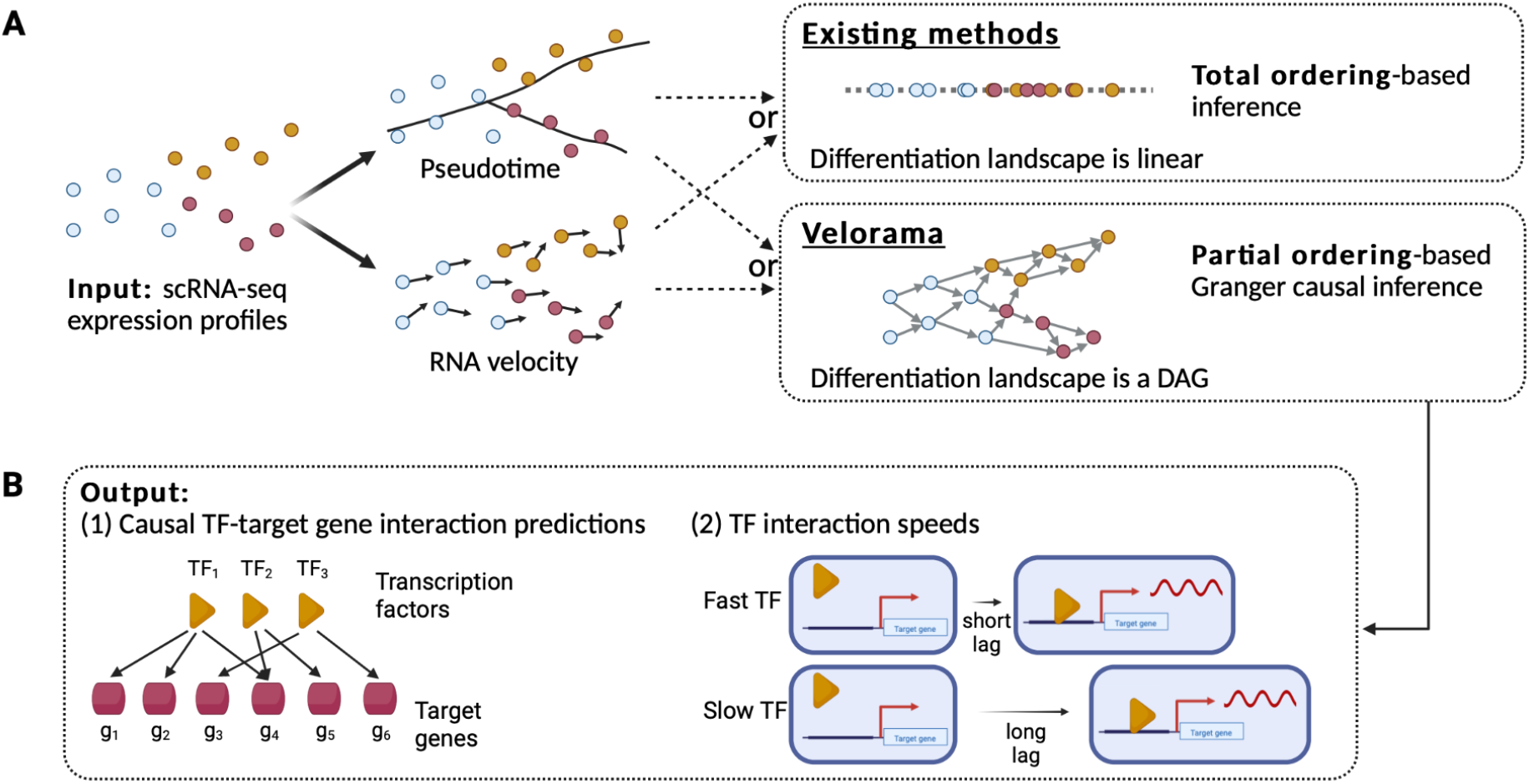
Velorama motivation and workflow. (A) Velorama takes as input scRNA-seq expression profiles to perform gene regulatory network (GRN) (GRN) inference. Existing methods need to impose a total (i.e., linear) ordering on cells in order to infer GRNs. Our key insight is that the differentiation landscape is more faithfully captured by a partial ordering, or directed acyclic graph (DAG), of cells constructed from RNA velocity or pseudotime. (B) Velorama uses a DAG of cells to predict causal interactions between transcription factors (TFs) and their target genes. Because Velorama focuses on Granger causal interactions that feature lagged interactions, it also predicts the interaction speeds for each TF–target gene interaction based on the lag between the TF’s expression change and the expression change of the downstream target gene.

A related and more pressing challenge is the integration of RNA velocity estimates into GRN inference methods. RNA velocity compares the spliced and unspliced counts of genes to predict per-cell state transitions during a dynamic biological process [12, 13]. It has proven powerful in understanding how differentiating cells commit to specific fates [14], and Qiu et al. [9] have argued that it more faithfully captures the expression dynamics of differentiation than pseudotime. These advantages suggest that the rich information encoded in RNA velocity data could be beneficial for inferring gene regulatory interactions. However, it is unclear how it can be combined with existing GRN approaches, as these approaches require a global pseudotemporal ordering over a population of cells while RNA velocity is an inherently local measure that estimates the state transitions for each cell. A potential workaround might involve using RNA velocity vectors to forcibly collapse cells into a global pseudotime-series over the full dataset, but doing so removes much of the rich local information provided by this data.

Here, we present Velorama, a novel method for gene regulatory network inference on scRNA-seq data that does not require a total ordering over the differentiation landscape. While some existing methods incorporate gene-wise RNA velocity for GRN inference [9, 15, 16], they do not account for how gene-wise velocities aggregate into cell state transitions. Velorama takes advantage of recent advances in cell fate mapping using RNA velocity [14], and is the first method to fully incorporate cell-to-cell transition probabilities for GRN inference. Our key conceptual advance is to perform causal GRN inference while modeling the differentiation landscape as a *partial*, rather than total, ordering of cells. Specifically, we model cell differentiation in a local, rather than global, context, which enables us to better capture cell dynamics. Instead of a linear pseudotemporal axis, we perform inference on a directed acyclic graph (DAG) of cells that encodes the partial ordering: edges in this graph connect transcriptionally related cells, with each directed edge capturing the direction of a cell’s local state transitions (**Figure 1A**).

To address the algorithmic challenge of performing causal inference on a partial ordering, we build upon recent innovations in Granger causal (GC) inference. In a gene regulatory context, Granger causality reasons that a causal regulatory gene’s past expression will be significantly predictive of its target gene’s current expression. While standard GC inference approaches are limited to total orderings, we recently introduced a generalization of Granger causality to partial orderings that uses a graph neural network framework to perform GC inference on a DAG [17]. Our previous work assessed pairwise interactions between each ATAC-seq peak (chromatin accessibility) and a candidate gene in its genomic neighborhood. However, this is insufficient for assessing gene–gene interactions, as transcriptional regulation often requires the coordination of multiple co-regulators [18–20]. We therefore introduce here a different mathematical formulation of GC inference that allows us to simultaneously consider *all* potential TF and their non-linear, combinatorial effects. In Velorama, TF–target gene comparisons are made locally for each node in the DAG by relating the target gene’s value at a node to the values of *all* candidate TFs at the node’s ancestors (**Figure 1B**).

We evaluated Velorama on a variety of synthetic and real scRNA-seq datasets, on which it demonstrated considerably greater accuracy than existing methods. On synthetic datasets with associated ground-truth GRNs made available by Dibaeinia et al. [21], we first applied Velorama using only pseudotime data, bench-marking it against other GRN methods. Velorama offers state-of-the-art precision and recall, substantially outperforming a diverse set of pseudotime-based GRN inference techniques on the area under precision recall curve (AUPRC) metric. By applying Velorama on RNA velocity data, generated from the same ground-truth GRNs, we show that the use of RNA velocity can dramatically improve the quality of GRN inference, leading to an additional 1.8-3x improvement in AUPRC over what was achieved with pseudotime. These findings suggest that Velorama can help unlock insights unattainable by current methods. When applied real datasets characterizing brain, pancreas, and bone marrow differentiation, RNA velocity-based inference captures relevant regulatory interactions comparably or better than pseudotime.

We also used Velorama to specifically investigate the regulatory dynamics of human corticogenesis, which uncovered intriguing evidence for the relationship between the speed of a TF’s interactions with its targets and its regulatory function. In particular, we observed that slow TFs were significantly linked to gliomas, were less cell type-specific, and co-regulated targets preferentially with faster TFs. On the other hand, fast TFs were implicated in neuropsychiatric disorders and were more cell type-specific. As RNA velocity assays are increasingly being leveraged to reveal fine-grained dynamics of differentiation and disease, Velorama’s unprecedented ability to identify causal regulatory drivers from RNA velocity data opens the door to uncovering deeper mechanistic insights into dynamic gene regulatory processes.

## 3 Results

### 3.1 Overview of Velorama

Velorama infers causal TF–target gene interactions for a single-cell dataset by representing the dataset’s expression dynamics as a DAG of cells. This DAG of cells can be constructed from pseudotime or RNA velocity measurements. When only pseudotime information is available, we construct the DAG by orienting the edges of a *k*-nearest-neighbor graph of cells in the direction of increasing pseudotime, which allows us to appropriately model merging and branching trajectories without collapsing them onto a single axis. When RNA velocity estimates are available, the DAG is constructed directly from the RNA velocity-based cell–cell transition probabilities inferred by single-cell fate mapping tools, like CellRank [14].

Granger causality is an econometric formulation to estimate temporal causality from a time series of multi-variate observations. It leverages the intuition that a cause precedes its effect and therefore, temporally-predictive relationships between dynamically changing variables could be indicative of causal relationships [22]. The inference procedure entails a statistical test to evaluate whether the lagged values of a candidate causal variable are predictive of the effect variable’s current value. A general approach to doing so uses regularization: the coefficients associated with the candidate causal variables are regularized, such that only the most relevant predictors are retained. With Velorama, we sought to perform Granger causal inference of TF–target gene interactions [17]. However, with classical Granger causality we run into the limitation that it requires a sequential ordering over which to infer lagged relationships.

Addressing this limitation is one of Velorama’s key conceptual advances: we represent lagged relationships as a DAG, with a node being “preceded” by its ancestors in the directed graph (**Figure 2**). Mathematically, DAGs encode partial orderings; these are a superset of total (i.e. sequential) orderings. With this motivation, we designed a new algorithm to perform Granger causal inference on a DAG of cells in order to compute the lagged expression of candidate TFs for a target gene. We consider these lagged TF expression values as predictors for the expression of the target gene in a feedforward neural network model (**Figure S1**) [23]. The first layer of the neural network is sparsified via regularization, enabling the weights associated with each TF at each lag point to be interpreted for Granger causality. As all TFs are considered simultaneously for each target gene, the inferred Granger causal interactions account for possible TF cooperativity. The final prediction for each candidate TF–target gene interaction is derived from an ensemble of neural network models, each of which is trained with a different sparsity penalty.

**Figure 2:**
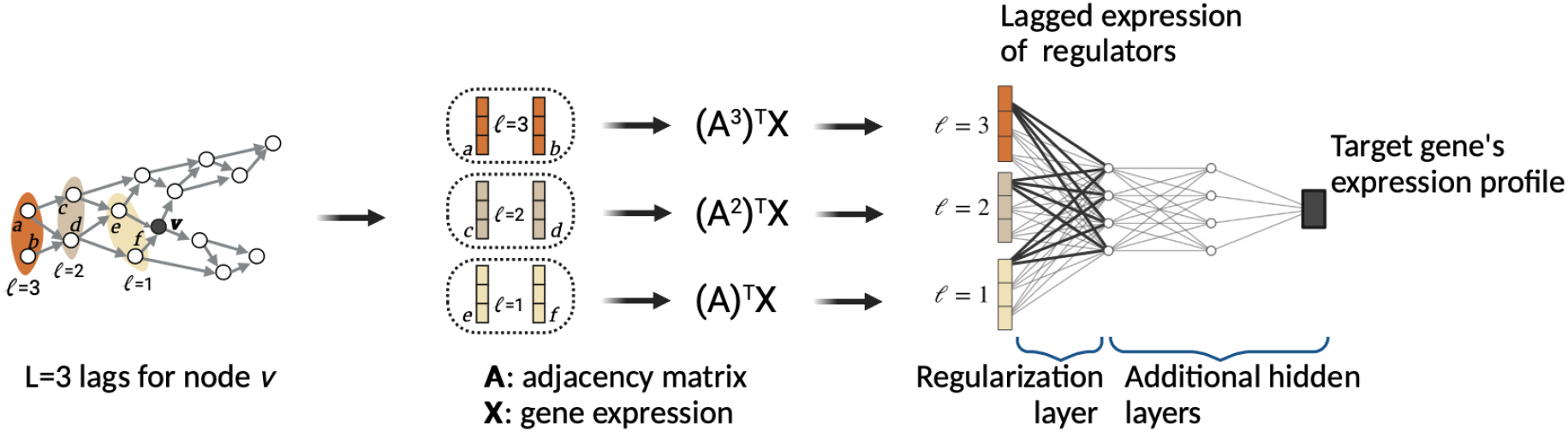
Velorama architecture. We perform Granger causal inference on a DAG of cells, taking advantage of a recent advance in generalizing Granger causality with the use of graph neural networks. At each node (cell), we collect the expression of putative TFs (e.g., three genes per cell) at the cell’s ancestor nodes (e.g., lag *L* = 3). These lagged regulator expression values are considered as putative predictors of the target gene’s expression. A key component of our neural network is the first hidden layer where regularization of the weights is applied to identify causal TF–target relationships.

The pseudotime setting of Velorama can operate with just the single-cell gene expression data and a root cell (for pseudotime computation) as the input, while the RNA velocity setting only requires spliced and unspliced transcript counts as input. The output is a ranked list of TF–target gene interactions along with the estimated speeds for each of these interactions. A more detailed description of the DAG construction procedure, Velorama architecture, and training process is presented in **STAR Methods**.

### 3.2 Velorama accurately infers GRNs from pseudotime-based expression dynamics

We benchmarked Velorama on a series of synthetic datasets for which the underlying ground-truth GRN is known. We used four differentiation datasets from SERGIO [21], which simulates single-cell expression dynamics according to a given GRN. These datasets, which we obtained from the SERGIO GitHub repository, span a range of differentiation landscapes, including linear (Dataset A), bifurcation (Dataset B), trifurcation (Dataset C), and tree-shaped (Dataset D) trajectories (**Figure 3A, STAR Methods**). The underlying GRN for each of these datasets consists of 100 genes and 137 edges. For each of these GRNs, SERGIO simulates spliced and unspliced counts as well as total transcript counts.

**Figure 3:**
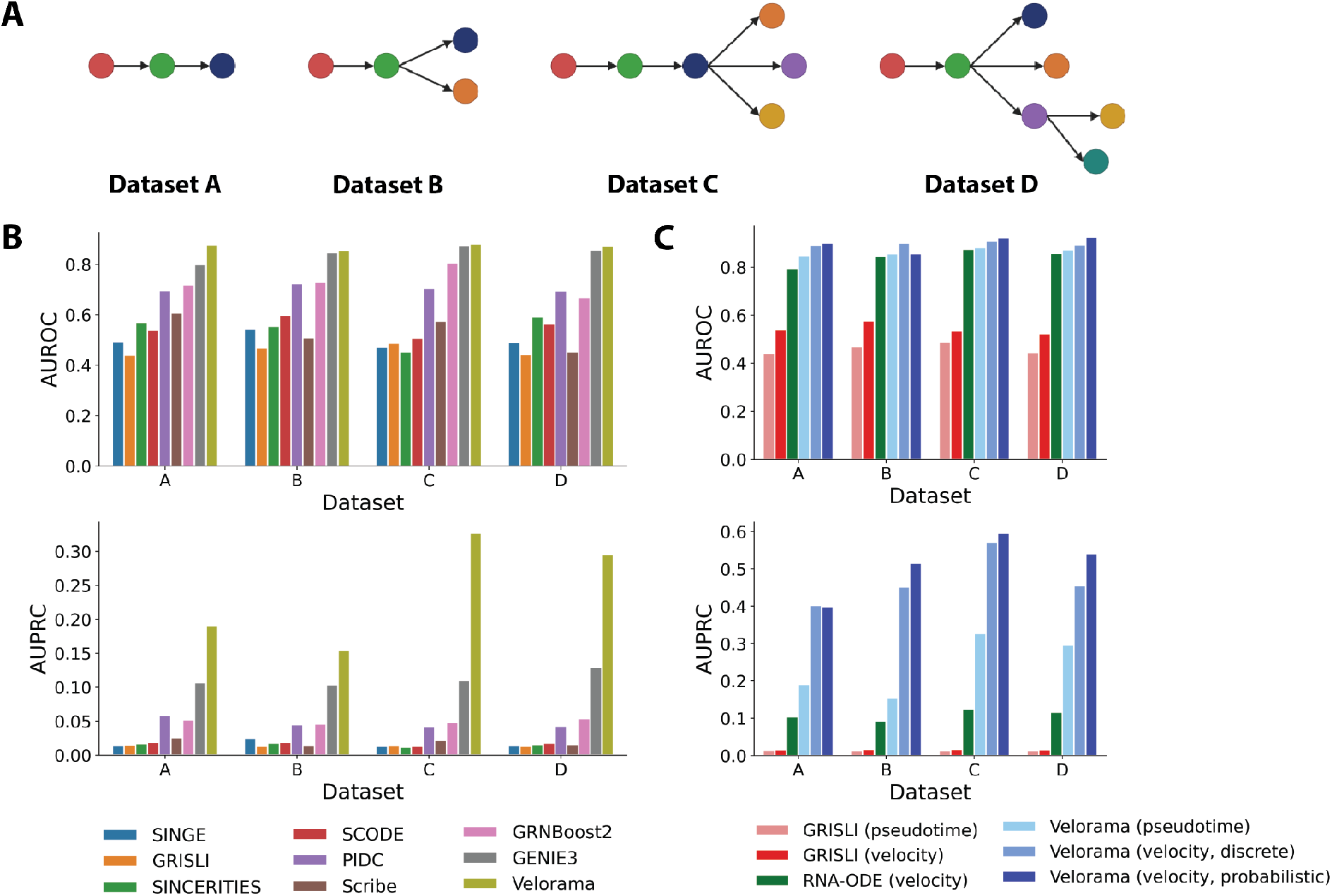
Evaluation of GRN Inference Accuracy for Single-Cell Differentiation Data. (A) Differentiation trajectories for the four synthetic datasets sourced from SERGIO [21]. Each node corresponds to a distinct cell state and directed edges represent cell state transitions along a trajectory. (B) Comparison of the accuracy of GRN inference algorithms in predicting the ground-truth network for each of the SERGIO synthetic datasets. Pseudotime data is used in this comparison for methods that require an ordering of cells. (C) GRN inference accuracy for Velorama across different settings for leveraging pseudotime or RNA velocity dynamics. In addition to Velorama, we show the performance of RNA-ODE and GRISLI, which also use RNA velocity for GRN inference.

We first sought to evaluate Velorama when only pseudotime data is available. Any benchmarking of Velorama requires this setting since existing methods can not make use of the full RNA velocity transition matrices. We compared against a diverse set of methods: GENIE3, GRNBoost2, GRISLI, Scribe, PIDC, SCODE, SINCERITIES, and SINGE [5–11, 15, 16]. We chose these methods because (a) their underlying mathematical and statistical techniques vary widely (decision trees, differential equations, information theory, and total ordering-based Granger causality), and (b) they were the top-rated methods in a comprehensive single-cell GRN benchmarking study [24].

On both the area under receiver operating curve (AUROC) and the area under precision-recall curve (AUPRC) metrics, Velorama outperforms all other methods (**Figure 3B**). Velorama’s substantial accuracy gains with respect to AUPRC are particularly notable, as the AUPRC metric has typically been the focus of GRN inference tasks due to the sparsity of GRNs and the consequent class imbalance associated with inferring its structure. On the area under receiver operating curve (AUROC) metric, Velorama also offers state-of-the-art performance, with it and GENIE3 being the two top-performing methods. GENIE3, originally designed for bulk RNA-seq, has consistently been reported to be one of the top-performing GRN methods. As such, Velorama’s outperformance against GENIE3 on the AUPRC metric and competitiveness on the AUROC metric is notable. The improvement of Velorama over SINGE and SINCERITIES, both of which utilize total ordering-based Granger causal inference, also suggests that our innovation of extending Granger causality to the DAG plays a crucial role in Velorama. We note that Velorama’s relative AUPRC outperformance is most substantial for Datasets C and D, which feature the most complex differentiation landscapes with the largest number of cell types and branching points in their respective trajectories (**Figure 3A**). These findings support the motivating intuition behind our work: compared to total orderings, partial orderings can more effectively capture a complex differentiation landscape.

#### Velorama flexibly incorporates time stamps from real time series datasets

Many single-cell experiments provide time stamp information [25], which we hypothesized could be incorporated into Velorama to better capture temporal relationships between cells and, hence, more accurate GRNs. We leveraged Schema [26] to synthesize the datasets’ time stamps with gene expression-based pseudotime estimates. We subsequently benchmarked Velorama using this approach on the SERGIO dataset and found that incorporating time stamps resulted in marked accuracy improvements for GRN inference (**STAR Methods, Table S1**).

We next analyzed two real single-cell time series datasets. The first profiled the progression of cardiac hypertrophy in cardiomyocytes [27], while the second profiled the response of A549 cells to pracinostat, an HDAC inhibitor [28]. In both cases, RNA velocity information was unavailable and we therefore applied Velorama with pseudotime to infer TF–target gene interactions. We identified the top 5% of all candidate TF–target gene interactions and evaluated the retained TFs for enrichment of GO terms related to the specific biological processes profiled in each dataset. For the cardiac hypertrophy dataset, we evaluated genes related to the regulation of cardiac muscle hypertrophy (GO:0010611) and focused on genes implicated in chromatin organization (GO:0006325) and remodeling (GO:0006338) for the A549 dataset given the impact of HDAC inhibitors on these processes. Even with no time stamp information, we observed a moderate enrichment for these GO terms (FDR *<* 0.1, Enrichr [29]). However, incorporating time stamps contributed to a stronger enrichment (**Table 1**). These results further highlight how Velorama can be flexibly used to unlock rich temporal information within single-cell data to reveal crucial regulatory interactions underlying disease and treatment-related processes.

**Table 1:** Enrichment of dataset-specific biological pathways for Velorama-inferred GRNs from single-cell time series data. Integrating time stamps alongside pseudotime with Velorama contributes to more significant enrichment of relevant pathways compared to Velorama with pseudotime alone.

| Method | cardiac hypertrophy | A549 + pracinostat |  |
| --- | --- | --- | --- |
|  | regulation of cardiac muscle hypertrophy | chromatin organization | chromatin remodeling |
| Velorama (pseudotime) | $2.2 \times 10^{-2}$ | $8.9 \times 10^{-2}$ | $7.7 \times 10^{-2}$ |
| Velorama (pseudotime + time stamps) | $9.6 \times 10^{-3}$ | $2.9 \times 10^{-5}$ | $5.4 \times 10^{-3}$ |

### 3.3 Velorama capitalizes on RNA velocity measurements

We also assessed Velorama’s ability to leverage RNA velocity for GRN inference when this data is readily accessible. To do so, we applied Velorama to the same synthetic datasets as above while taking advantage of the spliced and unspliced transcript counts provided by SERGIO. We weighed edges in the Velorama DAG according to the RNA velocity-based cell–cell transition probabilities estimated by CellRank and compared the GRN inference accuracy of this RNA velocity-based version of Velorama with that of the pseudotime-based version. We also performed an ablation on RNA velocity-based inference to test if Velorama could exploit the fine-grained information from the CellRank transition probabilities, rather than a binary edge-presence indicator. We refer to the setting which directly uses the CellRank transition probabilities as the “probabilistic” RNA velocity setting. In the ablated setting, which we refer to as the “discrete” setting, we retain only the edges in the CellRank transition matrix that correspond to the top 50% of each cell’s transition probabilities and binarize the resulting matrix prior to DAG construction.

Both RNA velocity settings of Velorama substantially improved upon the pseudotime version, even as the latter itself demonstrated state-of-the-art performance compared to existing benchmarks. Across the various datasets, the RNA velocity-based AUPRC metrics consistently improved upon the pseudotime-based metrics by 1.8–3x (**Figure 3C**). In addition, our ablation studies suggest that Velorama can exploit fine-grained cell transition probabilities derived from RNA velocity, with the probabilistic setting outperforming the discrete setting with respect to AUPRC for 3 of the 4 datasets. Indeed, with the probabilistic setting, Velorama’s improvement over existing methods is substantial, surpassing AUPRC values for GENIE3 predictions by 3–5x (**Figure 3B,C**). Velorama’s RNA velocity-based version also substantially outperforms existing RNA velocity-based approaches. For instance, RNA-ODE [16] uses random forests (like GENIE3) on RNA velocity data to infer GRNs. Its performance is similar to that of GENIE3, while Velorama’s AUPRC is nearly 4x higher. To confirm that these results were not a measurement artifact, we also visualized the actual ground-truth GRN matrix and the Velorama-inferred GRNs when using pseudotime or RNA velocity measurements (**Figure S2**). RNA velocity-based GRNs indeed capture the ground-truth regulatory relationships with higher fidelity.

The substantial performance gains associated with using RNA velocity in Velorama emphasize the value of accurately reconstructing cell-state transitions from scRNA-seq snapshots. Our results corroborate previous findings that RNA velocity better captures temporal couplings between single-cell measurements compared to pseudotime, thereby enabling more accurate causal GRN inference [9]. For instance, GRISLI’s RNA velocity-based version also outperforms its pseudotime-based version.

As with pseudotime, Velorama can incorporate user-provided time stamps from a time-course study with RNA velocity data to better estimate temporal relationships between cells (**STAR Methods**), driving additional improvements in GRN inference (**Table S1**). Altogether, these results point to the flexibility of Velorama to make full use of the rich temporal information revealed by scRNA-seq data to unravel regulatory drivers of complex differentiation landscapes.

### 3.4 Velorama identifies relevant regulatory interactions in real RNA velocity datasets

We applied Velorama to analyze gene regulatory interactions related to three scRNA-seq differentiation datasets. These datasets characterize the expression dynamics for mouse endocrinogenesis [30], mouse dentate gyrus neurogenesis [31], and human hematopoiesis [32] (**Figure 4A**). After performing standard pre-processing (**STAR Methods**), we applied Velorama with probabilistic RNA velocity, to infer TF–target gene interactions for each dataset. We separately also computed a diffusion pseudotime estimate for these datasets and applied Velorama with pseudotime. For each dataset, we first identified the top 5% of all candidate TF–target gene interactions based on the same scoring scheme as above. The TFs implicated in these interactions were significantly enriched for Gene Ontology (GO) terms corresponding to the specific biological processes being profiled in each of these datasets (FDR *<* 0.05, Enrichr [29], **Figure 4B**).

**Figure 4:**
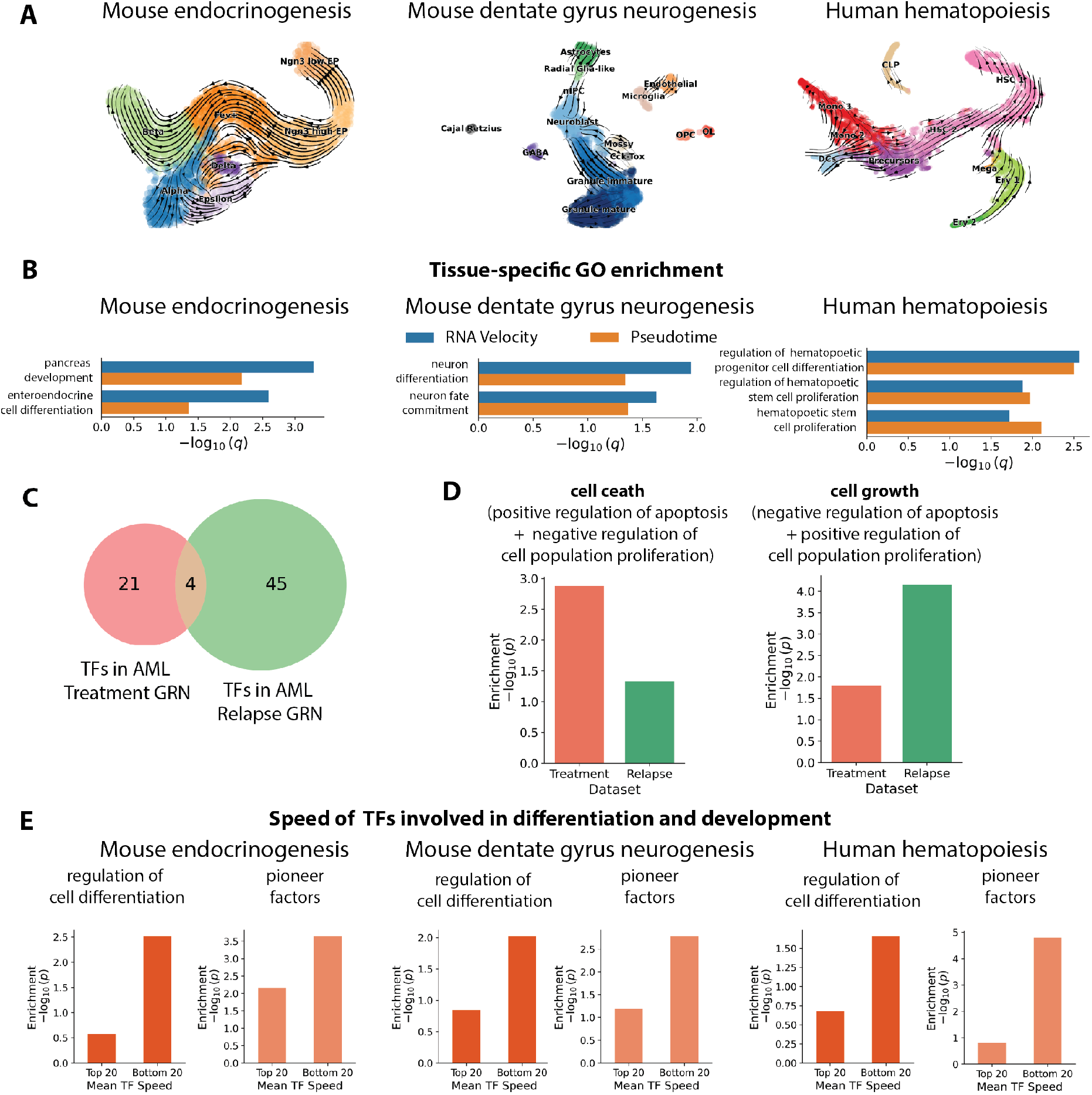
Velorama applications for studying regulatory dynamics on scRNA-seq RNA velocity datasets. (A) Stream plot visualizations of RNA velocity-based expression dynamics. (B) Enrichment of dataset-specific biological pathways for TFs identified by Velorama. Compared to pseudotime-based inference, RNA velocity-based inference is more strongly enriched for TFs known to be involved in relevant pathways. (C) Venn diagram of TFs implicated in the AML treatment GRN and the AML relapse GRN. (D) Enrichment of cell death-and cell growth-related gene programs for TFs in the AML treatment GRN and the AML relapse GRN. AML treatment is governed by TFs that are more significantly associated with cell death, while AML relapse is governed by TFs that are more significantly associated with cell growth. (E) Enrichment of differentiation GO terms and pioneer factors for the 20 fastest and slowest TFs identified by Velorama. Differentiation-related TFs and pioneer factors are associated with slower interaction speeds.

We also validated the causal direction of the Velorama-inferred TF–target gene interactions by examining them against the true time stamps available in time-series data. We found that Velorama GRNs were indeed enriched for TF–target gene interactions that align with the temporal flow of causal gene regulation, in which TF activation precedes the differential expression of its downstream target (**Table S2**). In addition, we compared Velorama with scDiff [25], which explicitly uses time stamps to infer TF–target gene interactions. We applied Velorama solely using RNA velocity, without using the data’s time stamps, and nonetheless found substantial overlap in the TF–target gene interactions inferred by Velorama and scDiff (**Figure S3**). This result suggests that RNA velocity’s can help recover true temporal information from single-cell data without the direct utilization of time stamps (**Figure S4**) [9] and further demonstrates the power of leveraging RNA velocity for causal GRN inference.

Moreover, when comparing this enrichment for interactions inferred when pseudotime instead of RNA velocity was supplied to Velorama, the RNA velocity-based interactions yielded more significant enrichment for 5 of the 7 GO terms. This improvement in enrichment is possibly a consequence of the complex expression dynamics of these datasets, comprising multiple cell types and differentiation branches, which may be challenging for previous pseudotime-based approaches that collapse these dynamics onto a total ordering of cells. Notably, even with pseudotime, employing Velorama’s partial ordering-based representation helps alleviate these issues, as evidenced by the significant enrichment results for that setting. Nonetheless, the RNA velocity-based DAG better handles these dynamics.

To investigate the utility of Velorama and RNA velocity in studying complex diseases, we also analyzed two acute myeloid leukemia (AML) datasets for which both spliced and unspliced transcript counts were available. The two datasets enabled a relapse–vs.–treatment comparison: the first dataset profiled AML relapse [33], while the other profiled AML cells responding to a combination treatment of venetoclax and azacitidine [34]. We used Velorama to infer a GRN of the highest-scoring 5% of TF–target gene interactions for each of the AML relapse and treatment datasets. We then examined the TFs in each of these GRNs and identified substantial differences in the regulatory drivers that distinguish AML progression and effective treatment (**Figure 4C**). We observed that the TFs in the AML relapse GRN were enriched for key genes implicated in the negative regulation of apoptosis and positive regulation of cell proliferation, including TOX and XBP1. On the other hand, the TFs in the treatment GRN were enriched for genes that drive the positive regulation of apoptosis and negative regulation cell proliferation, such as FOXO3, RUNX3, and IRF1. (**Figure 4D**). These enriched biological processes cohere with the regulatory mechanisms that are expected to underlie uncontrolled cell growth in cancer progression and cell death in effective cancer treatment [35, 36]. Taken together, these results underscore that RNA velocity-driven GRN inference with Velorama can be effective in detecting disease-relevant gene dysregulation.

### 3.5 Velorama suggests certain classes of transcription factors act more slowly than others

Velorama’s ability to estimate the relevant lags for Granger causal interactions enables it to predict the speed of each regulator–target gene interaction (**STAR Methods**). We calculated the interaction speeds for the top 5% of TF–target gene interactions in each of the three differentiation datasets. For each of the datasets, the bottom 20 TFs by interaction speed were significantly enriched for GO terms related to cellular differentiation (GO:0045595) (*p <* 0.05, hypergeometric test, **Figure 4E**). In comparison, the 20 regulators with the highest interaction speeds did not show enrichment for these terms. This pattern of enrichment for only the lower interaction speed regulators was observed not just for each tissue but across all three tissues.

This observation aligns with known mechanisms of gene regulatory control during differentiation. Regulators of these processes often participate in a number of epigenetic and/or cofactor recruitment steps prior to influencing their target [37, 38]. As a result, these intermediary regulatory activities may manifest in a lag between changes in TFs’ expression levels and those of their targets, hence leading to lower observed interaction speeds for such TFs. To explore this phenomenon further, we evaluated the relationship between the speed of TFs and their ability to act as pioneer factors. Pioneer factors have the ability to access closed chromatin, facilitate chromatin opening, and mediate binding of other TFs, making them critical regulators for which we expect to observe a lag between their initial expression and their downstream effects [39]. Using a curated list of TFs with pioneer-like properties [40], we found that the 20 slowest TFs by interaction speed were significantly enriched for these pioneer factors across all three differentiation datasets (*p <* 0.05, hypergeometric test, **Figure 4E**). Meanwhile, enrichment for these pioneer factors in the 20 fastest TFs was only observed in the mouse endocrinogenesis dataset. Identifying the regulatory range and duration of transcription factors is a major open problem and this preliminary investigation suggests a promising direction of future work: RNA velocity measurements, analyzed with Velorama, could systematically measure regulatory speed in a variety of tissues, leading to a deeper understanding of gene regulatory control.

### 3.6 Velorama reveals functional and disease distinctions between fast and slow TFs in corticogenesis

The development of the human cortex requires the precise coordination of numerous regulatory processes [41]. To determine the role of temporal coordination of human corticogenesis, we evaluated a single-cell multimodal dataset profiling human fetal cortical samples during mid-gestation [42]. This dataset includes single-cell gene expression and chromatin accessibility profiles derived from the same cell. We applied Velorama to the gene expression component of this dataset to construct a map of causal TF–target interactions for cells along the excitatory neuron differentiation (**Figure 5A, STAR Methods**). We focus on the top 1% of all candidate TF–target interactions. The TFs retained in these interactions are significantly enriched for genes implicated in neuron generation and neuron differentiation (FDR *<* 0.001, Enrichr [29]).

**Figure 5:**
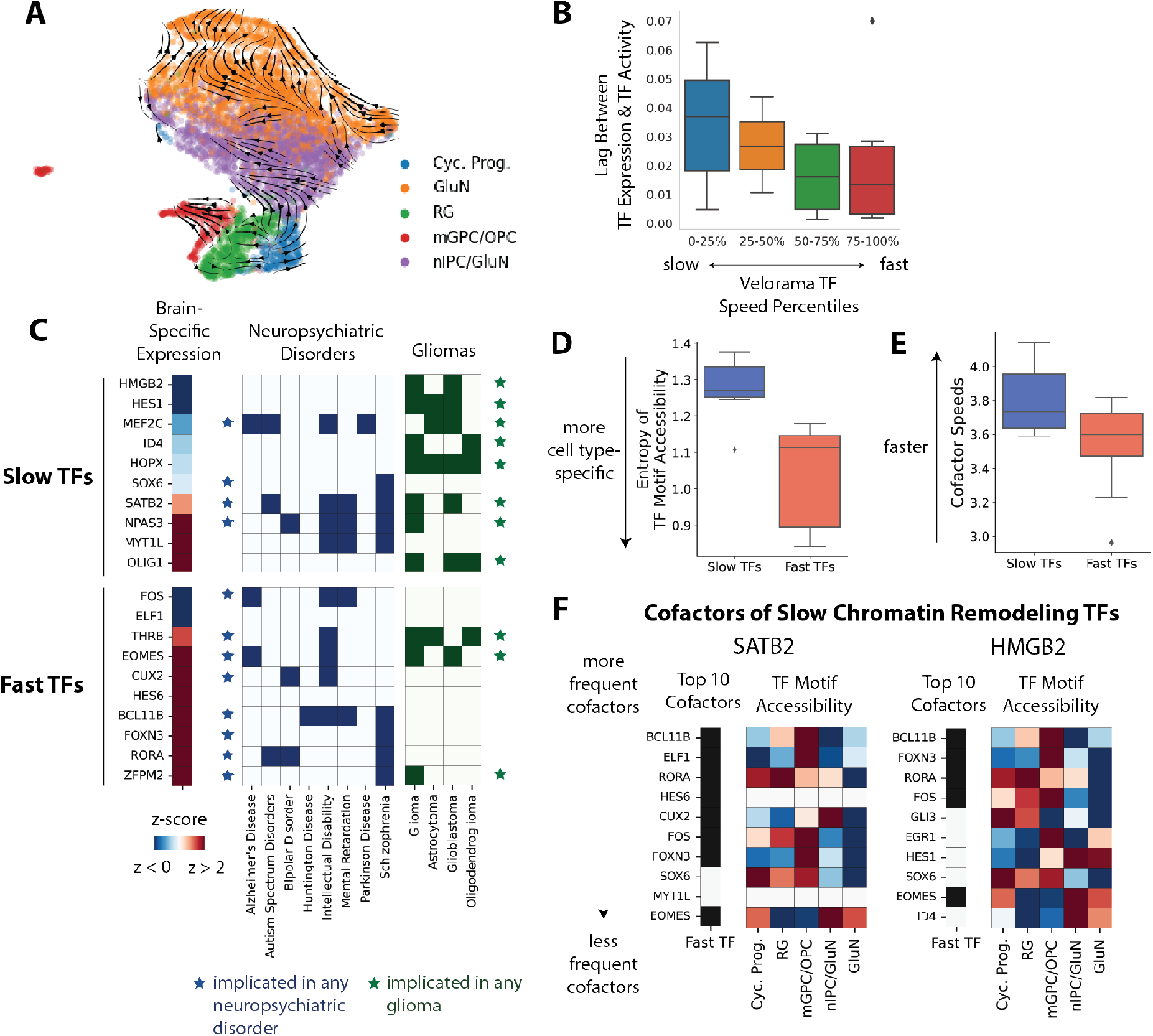
Velorama enables the dissection of temporal regulatory control and coordination in human corticogenesis. (A) Stream plot visualization of RNA velocity-based expression dynamics for corticogenesis. (B) Velorama-inferred TF speeds cohere with estimated lags between TF expression and TF activity. The activity of a TF is derived from the accessibility of chromatin regions containing the TF’s motif. (C) Brain-specific expression levels and disease relevance for “slow TFs” and “fast TFs”. (D) Comparison of the cell type-specificity of motif accessibility for “slow TFs” versus “fast TFs”. (E) Comparison of the cofactor speeds for “slow TFs” versus “fast TFs”. (F) Cofactors of the “slow TFs” SATB2 and HMGB2, both of which have chromatin remodeling functions. Heatmaps represent (left) whether a cofactor was identified as a “fast TFs” and (right) the mean TF motif accessibility in each of the cell types in the excitatory neuron trajectory.

We also evaluated Velorama ‘s TF interaction speed estimates by jointly leveraging the dataset’s single-cell gene expression and chromatin accessibility profiles. Using the chromatin accessibility data, we first used chromVAR [43] to infer TF activities in each cell based on the presence of TF motifs within accessible regions of the genome. For each TF, we then computed the lag between its gene expression and its corresponding TF motif activity (**STAR Methods**). We found that this TF expression-activity lag was anti-correlated with Velorama-inferred TF interaction speeds, with the slowest TFs inferred by Velorama associated with the longest lags (**Figure 5B, S5**). These results, therefore, verify that Velorama can accurately predict not only the causal links between TFs and their target genes but also the time delays that are implicated in their causal interactions. Its ability to do so, therefore, opens the door for uncovering insights about the relationship between gene regulatory timing and function.

To understand the role of temporal control in corticogenesis, we analyzed the TFs with the slowest and fastest interaction speeds. We refer to the 10 fastest and slowest TFs as the “fast TFs” and “slow TFs”, respectively. We found that 7 of the 10 “fast TFs” have highly brain-specific expression levels (*z*-score *>* 2, **STAR Methods**), while only 3 of the “slow TFs” are brain-specific (**Figure 5C**), indicating potentially distinct functions for these two categories of TFs.

We first investigated the disease relevance of these TFs by analyzing their annotations in DisGeNET [44]. We observed that the “fast TFs” are significantly enriched for gene sets implicated in neuropsychiatric disorders (7 of 10 TFs; *p <* 0.001, hypergeometric test), while the “slow TFs” do not display this degree of enrichment (5 of 10 TFs; *p >* 0.05, hypergeometric test). On the other hand, “slow TFs” appear to be highly enriched for gene sets implicated in gliomas (8 of 10; *p <* 10^−5^, hypergeometric test), while “fast TFs” are not as relevant in gliomas (2 of 10; *p >* 0.1). These “slow TFs” are also significantly associated with more general regulatory processes, including chromatin organization and development processes linked to other non-brain tissue types, such as cartilage development and cardiac muscle differentiation (*p <* 0.01, hypergeometric test). Taken together, these results point to the relative importance of “slow TFs” in more universal regulatory functions that, when disrupted, may contribute to broad phenotypic consequences like cancer. Meanwhile, the stronger link between “fast TFs” and neuropsychiatric diseases potentially indicates more precise, neural-specific roles for these TFs in neural development and maintenance.

Based on these observations, we then hypothesized that the more specific functions of “fast TFs” would be reflected in the cell type-specificity of the chromatin regions on which they bind. To test this, for each TF we evaluated the accessibility of chromatin regions containing its motif and compared this accessibility across the different cell types profiled along the corticogenesis trajectory. We indeed found that the “fast TFs” are associated with significantly more cell type-specific chromatin accessibility patterns at loci containing their motifs than “slow TFs” (*p <* 0.01, *t*-test; **Figure 5D, STAR Methods**). This result, therefore, provides further evidence for the disparities between not only the specificity of the functions of “slow TFs” and “fast TFs” but also the mechanisms by which they affect gene regulation.

These differences between the “slow TFs” and “fast TFs” led us to explore how TFs of varying interaction speeds might work cooperatively to regulate gene expression. In particular, we took advantage of Velorama ‘s ability to account for the combinatorial effects of TFs to identify cofactors of “slow TFs” and “fast TFs”. The set of top cofactors for a particular TF was defined to be the 10 TFs that most frequently co-occur with that TF upon accounting for the abundance of targets per TF (**STAR Methods**). We found that the interaction speeds for the cofactors of “slow TFs” were significantly higher than those of the “fast TFs” (*p <* 0.05, one-tailed *t*-test; **Figure 5E**). This finding indicates a mechanism for gene regulatory control in which TFs work cooperatively with cofactors that have interaction speeds different from their own. Based on the functional and cell type-specific differences between TFs of different interaction speeds described above, this result indicates that combinations of TFs with distinct interaction speeds and distinct functions are implicated in co-regulatory processes.

To explore the cooperativity of “slow TFs” and “fast TFs” more specifically, we examined the cofactors of two “slow TFs” with known chromatin remodeling capabilities -SATB2 and HMGB2. SATB2 is known to remodel chromatin in differentiating cortical neurons, and HMGB2 can bend DNA to facilitate cooperative interactions between cis-acting TFs [45, 46]. Each of these TFs are linked to many cofactors that act as “fast TFs”, with 8 of the top 10 most frequently co-occurring cofactors for SATB2 found to be “fast TFs” and 5 of the top 10 for HMGB2 (**Figure 5F**). Of these “fast TF” cofactors, BCL11B and EOMES were among the cofactors linked to both SATB2 and HMGB2. Intriguingly, BCL11B and EOMES have also both been previously shown to co-regulate gene expression with SATB2 to drive cell fate determination. Specifically, BCL11B is known to regulate glial progenitor cell and oligodendrocyte proliferation, and EOMES is known to be a major determinant of intermediate neural progenitors [47–49]. This cell type-specific relevance for BCL11B and EOMES corresponds exactly to the cell type-specificity of accessible chromatin possessing their motifs, with chromatin regions for BCL11B and EOMES motifs being most accessible in the multipotent glial/oligodendrocyte progenitor (mGPC/OPC) and intermediate neural progenitor (nIPC/GluN1) cell type categories, respectively (**Figure 5F**). As such, these results suggest the possibility of “slow TFs”, like the SATB2 and HMGB2 chromatin remodelers, working in coordination with “fast TFs”, like BCL11B and EOMES, to drive cell type-specific functions and cell fate specification in differentiation. The ability of Velorama to capture the dynamics of TF activity as well as the interplay of these dynamics with TF cooperativity underscores its unique potential for unlocking key insights about causal regulatory dynamics.

## 4 Discussion

scRNA-seq datasets capture rich information about gene regulatory dynamics, parts of which are neglected by current techniques due to their inability to operate on partial, nonlinear orderings of cells. This is particularly a problem for RNA velocity data, since many existing methods require a pseudotime estimate. An alternative approach is described in Scribe [9] where RNA velocity is considered, in essence, as two time-points of scRNA-seq data: one corresponding to spliced counts (regulators) and the other to unspliced counts (targets). However, this approach makes the heavily-questioned assumption that splice rates across genes are equal [13], is not designed to identify interactions spanning multiple cell states, and is unable to take advantage of cell transition estimates offered by methods like scVelo and CellRank [14, 50]. Velorama takes a conceptually different approach by modeling the differentiation dynamics suggested by RNA velocity as a DAG. Velorama is also compatible with pseudotime if RNA velocity is unavailable. While our previous work introduced DAG-based representations of cellular dynamics [17], Velorama is the first to integrate per-cell transition probabilities from RNA velocity for GRN inference. Velorama is also the first method to leverage these DAG-based dynamics to evaluate multiple candidate regulators simultaneously along with their combinatorial effects.

Velorama’s effectiveness on the GRN inference task opens the door to several directions for future work. Firstly, while Velorama has demonstrated notable performance using only scRNA-seq data, integrating this data with other data modalities, like chromatin accessibility and/or TF binding data, may contribute to further performance gains [26]. In addition, In addition, we have demonstrated the effectiveness of using Velorama with either pseudotime or RNA velocity, but opportunities exist to integrate these two approaches. By ensembling cell-cell similarity predictions from these two approaches, we can potentially harness the unique features captured by both to more faithfully represent the underlying cellular dynamics used by Velorama for GRN inference. Lastly, Velorama’s unique ability to infer the speeds of TF–target gene interactions newly enables the discovery of crucial links between temporal control of gene expression, its role in molecular functions, and disease etiology. Additional investigations of TF interaction speeds from the vast compendium of existing single-cell data can, therefore, further our understanding of time as a critical axis of gene regulatory control.

Velorama unlocks crucial information from RNA velocity that otherwise would be lost with other techniques. As efforts to model gene expression dynamics from RNA velocity continue to grow and improve, Velorama’s distinctive ability to flexibly leverage RNA velocity for causal GRN inference positions it to be a valuable tool for uncovering novel mechanistic insights into differentiation and disease.

## 5 STAR Methods

### 5.1 Key Resources Table

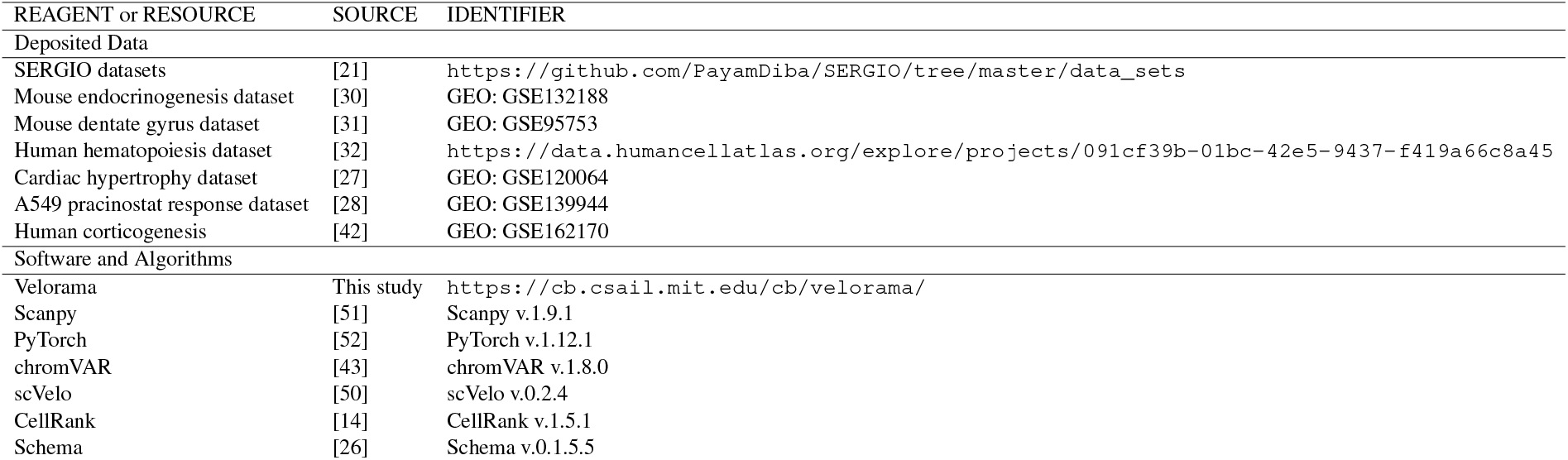

### 5.2 Method Details

#### 5.2.1 Classical Granger causal inference and its non-linear extensions

Granger causal (GC) inference, a statistical approach for estimating causal relationships within dynamical systems, has been effectively used in many settings for which only observational data is available [53–57]. Granger causality leverages the key intuition that since a cause precedes its effect, temporally-predictive relationships between dynamically changing variables could be indicative of causal relationships.

Statistical and machine learning methods have been developed to recover such temporally-predictive relationships. Broadly, there are two strategies to assess the existence of a GC relationship between variables *x* and *y*: ablation and invariance [17]. The ablation approach involves training two predictive models of *y*, denoted as *u* and *u*_*\x*_, where *u* includes the history of every variable in the system and *u*_*\x*_ excludes the history of *x*. If *u* performs significantly better than *u*_*\x*_, as determined by a one-tailed F-test, then *x* Granger-causes *y*. The invariance approach involves training just one predictive model of *y*, denoted as *f*, using the history of every variable. In this approach, *x* does *not* Granger-cause *y* if and only if the learned weights governing the interaction between *x* and *y* are all equal to 0. Equivalently, no causal relationship exists exactly when the prediction of *y* is invariant to the history of *x*. Unlike our previous work [17], here we chose the invarance-based approach since it allows multiple candidate Granger causal variables to be evaluated simultaneously, enables the lag associated with each Granger causal interaction to be determined, and reduces training time.

While the classic formulation of GC inference assumes linear interactions, extensions have been developed to capture the nonlinear interactions more often seen in real-world datasets. Many of these extensions have been based on deep learning-based architectures. Tank et al. [58] introduce a regularized multilayer perceptron and a long short-term memory network that model nonlinear relationships while simultaneously determining the lag of each putative causal relationship. Marcinkevičs and Vogt [59] extend self-explaining neural networks to multivariate time series data in order to determine whether a causal relationship induces a positive or negative effect.

Given a dynamical system with a totally-ordered sequence of *N* observations, each over *G* variables, Granger causal inference involves training per-variable models *f*_1_, *f*_2_, …, *f*_*G*_. Here, *f*_*j*_ models variable *j* as a function of the previous *L* observations:

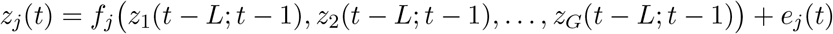

Here, *z*_*j*_(*t*) denotes the value of the variable *j* at observation *t, e*_*j*_(*t*) denotes an error term, and *z*_*k*_(*t* −*L*; *t* −1) is the sequence of *L* past observations of variable *k*: (*z*_*k*_(*t* − *L*), …, *z*_*k*_(*t* − 1)) [58]. Each pair of variables (*i, j*) has an associated weight tensor **W**_(*i,j*)_ that defines how variable *j* depends on past lags of variable *i*. The sub-tensor 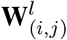 refers to the lag-*l* value of interaction between *z*_*i*_(*t* − *l*) and *z*_*j*_(*t*). Variable *i* is then said to Granger-cause *j* if |**W**_(*i,j*)_ | ≠ 0, meaning that *f*_*j*_ is not invariant to *z*_*i*_. During training, regularization can be applied to each weight tensor **W**_(*i,j*)_ to assist in achieving exact zeros for the non-causal relationships. By applying additional regularization terms on each 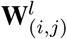, we can automatically detect the relevant lags of each putative causal relationship [58].

#### 5.2.2 Velorama: non-linear Granger causal inference on partially ordered observations

In both the linear and non-linear formulations above, *z*_*t*_ is defined to temporally precede *z*_*t*+1_, which implies the requirement of a total ordering on the *N* observations. Thus, dynamical systems that consist of branching points, such as cellular differentiation trajectories, are not compatible with these approaches. To address this, Velorama extends the nonlinearity and automatic lag selection of modern GC methods to DAG-structured dynamical systems that encode a partial ordering of observations.

Velorama takes in two inputs, **A** ∈ **ℝ**^***N****×* ***N***^ and **X** ∈ **ℝ**^***N****×* ***G***^, and produces one output **GC** ∈ **ℝ**^***G****×* ***G***^. **A** is the adjacency matrix of the DAG, where each node represents an observation and edges connect these observations. **X** is the feature matrix which describes the values of the *G* variables over the *N* observations, and **GC** summarizes the inferred causal graph, where **GC**_*ij*_ represents the strength of the causal relationship between variables *i* and *j*. We pre-compute a modified matrix **A**^*′*^, defined to be the normalized transpose of **A**, where row sums to 1 so as to account for variability in the in-degrees of the DAG. If a row of **A**^*T*^ consists of all zeros, the row is unaltered. **A**^*′*^ serves as a graph diffusion operator that aggregates information over each observation’s ancestors.

To infer Granger causal relationships, we train *G* separate models *f*_1_, *f*_2_, …, *f*_*G*_, each of which has the same architecture. We propose *f*_*j*_ to be a multilayer neural network that models variable *j* as a nonlinear function of ancestors within the DAG. The key component of our model architecture is the first hidden layer, which takes on the form

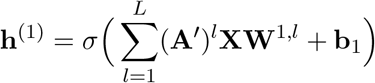

Here, (**A**′)^*l*^ represents the *l*^th^ power of **A**′, **W**^1,*l*^ ∈ **ℝ**^***G****×* ***d***^ is a learned weight matrix and **b**_1_ ∈ **ℝ**^***N****×* ***d***^ is a learned bias term. *d* is the number of hidden units per layer. (**A**′)^*l*^**XW**^1,*l*^ represents the information aggregated over *l* hops backward in the DAG (**Figure** S1a). Note that, as the **A**′ and **X** matrices are fixed, we can pre-compute (**A**′)^*l*^**X** = **A**′ (**A**′)^*l*−1^**X** inductively for 1 ≤ *l* ≤ *L*. *σ*(·) is a nonlinear activation function. In a *K* layer model, the hidden layers **h**^(*k*)^ for 2 ≤ *k < K* are given by

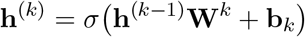

where **h**^(*k*)^ ∈ **ℝ**^***N****×* ***d***^, **W**^*k*^ ∈ **ℝ**^***d****×* ***d***^ and **b**_*k*_ ∈ **ℝ**^***N****×* ***d***^ (**Figure** S1b). Finally, the output *f*_*j*_ ∈ **ℝ**^***N***^ of the autoregressive model is given by

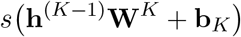

where **W**^*K*^ ∈ **ℝ**^***d****×****1***^ and **b**_*K*_ ∈ **ℝ**^***N****×* ***1***^. *s*(·) is an optional element-wise decoder function that links the real numbers to a domain-specific output.

##### Granger causal inference via regularization of W^1^

To infer causality with respect to variable *j*, we concatenate the per-lag matrices **W**^1,*l*^ from model *f*_*j*_ into **W**^1^ ∈ **ℝ**^***G****×* ***d***^. Let 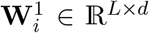 denote the weights that govern the interaction between variable *i* (TF) and variable *j* (target) (**Figure** S1c). Variable *i* does *not* Granger-cause variable *j* if 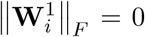, where ∥·∥_*F*_ denotes the Frobenius matrix norm. Similarly, we can infer these causal interactions with respect to a specific lag point as well, thereby enabling automatic lag selection in our framework. Let 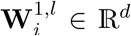 denote the weights that govern the interaction between variable *i* and variable *j* at a particular lag *l*. If variable *i* Granger-causes *j*, then *l* is not a relevant lag when 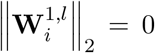, **Where** ∥·∥_2_ denotes the *ℓ*_2_ norm. We represent the Granger causal relationship between a candidate causal variable *i* and its target variable *j* at lag *l* as 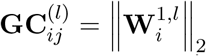.

To induce a sparse set of causal interactions, we simultaneously reduce the prediction error of each model *f*_*j*_ while encouraging the presence of exact zeros in the **W**^1^ matrices. Therefore, we define our loss function ℒ to be the sum of the mean-squared error loss *ℓ* (·,·) over all *j* models and a regularization penalty applied to the **W**^1^ matrices over all models:

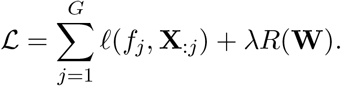

Here, **X**_:*j*_ ∈**ℝ**^***N***^ represents the value of variable *j* over all *N* observations. Let **W**^(*j*)^ denote the **W**^1^ corresponding to model *f*_*j*_ and **W** = [**W**^(1)^, …, **W**^(*G*)^] ∈ ℝ^*G*×*L*×*G*×*d*^ represent the concatenation of these matrices over all *G* models. *R*: ℝ^*G×L×G×d*^ → ℝ is the regularization loss function. The hyperparameter *λ* controls the regularization strength.

The specific form of *R* is guided by the regularization characteristics desired by the user. Tank et al. [58] introduced the hierarchical regularization scheme, which we use in our implementation. Hierarchical regularization penalizes longer lags more heavily than shorter lags, which has been found to achieve robustness with respect to the choice of the maximum lag *L* [58] (**Figure** S1d).

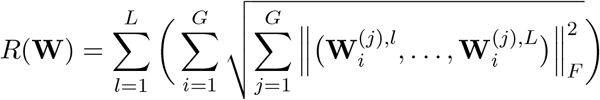

Traditional gradient descent algorithms such as Adam [60] and stochastic gradient descent often fail to converge the learned weights to exact zeros, hindering causal detection. We thus optimize the objective function ℒ via proximal gradient descent, a specialized algorithm designed to induce sparsity [61].

#### 5.2.3 Constructing the DAG from pseudotime or RNA velocity data

We first learn low-dimensional representations for each cell’s gene expression profile by performing principal component analysis (PCA) and projecting the gene expression profiles of cells onto the top *M* principal components. Optionally, if time stamps or additional data modalities are available (e.g., scRNA-seq time series or multimodal single-cell data), users can first endow the learned representations with this additional information by using multimodal integration techniques like Schema [26] (see below). We then construct a *k*-nearest-neighbor graph 𝒢 of cells based on Euclidean distances in the representation space. When using Velorama with pseudotime, we orient each edge *e* ∈ 𝒢 in the direction of increasing pseudotime. Doing so preserves the underlying differentiation structure while ensuring that the constructed graph is acyclic. For all datasets, we constructed the Velorama DAGs using *k* = 15 nearest neighbors and *M* = 50 principal components; Velorama’s effectiveness is robust to these hyperparameter choices (**Table S3,S4,S5,S6**).

When using Velorama with RNA velocity, we first apply CellRank to calculate cell–cell transition probabilities [14]. The probability *p*_*ij*_ of cell *i* transitioning into a neighboring cell *j* is calculated based on the alignment between the velocity vector of cell *i* and the displacement vector between cell *i*’s and *j*’s gene expression profiles. Consequently, these transition probabilities characterize cellular dynamics in a cell-specific, local manner that is conducive for representing cell state changes on single-cell graphs. As such, we directly use these transition probabilities as edge weights for the Velorama adjacency matrix **A**.

A possible complication with directly using the CellRank cell–cell transition matrix in Velorama is that this matrix may contain cycles. In Velorama, we only consider ancestors up to *L* hops away, so only cycles of up to length *L* are relevant. To combat the possibility of cycles in these *L* hops, we first remove cycles of length 2 by setting *p*_*ji*_ = 0 if *p*_*ij*_ *> p*_*ji*_. When we pre-compute powers of **A**′, we then set all non-zero diagonal terms back to zero after each successive multiplication with **A**′, as cycles can be identified by non-zero terms along the diagonal.

#### 5.2.4 Identification of Granger-causal interactions

The hyperparameter *λ* plays a key role in regularization, with higher values of *λ* leading to sparser GRNs. We generate an ensemble of models, trained over a range of *λ* values, and we compute the GRN by aggregating the models’ results across the ensemble. Thus, the user can simply use Velorama with its default settings. We sweep *λ* uniformly over the log-scaled range of [0.01, 10]. We retain only the per-lag **GC** ∈ **ℝ**^***G****×* ***G***^ matrices with between 1% and 95% non-zero values. To aggregate results over the various *λ* values, we then sum the retained **GC** matrices corresponding to each lag, which results in *L* composite **GC** matrices. These per-lag matrices are themselves summed to produce a single **GC** matrix, which we report as the GRN.

#### 5.2.5 TF–target gene interaction speed predictions

Velorama’s ability to perform lag selection enables it to predict the interaction speed for each TF–target gene pair. For a target gene *j*, we quantify the Velorama interaction speed between a TF *i* and the target by first scoring each of its associated lags *l* as the *ℓ*_2_-norm of the weights 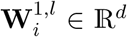. For each TF–target gene pair (*i, j*), we then identify the largest lag with a non-zero score. We average these largest lags over the ensemble of models to yield an estimated lag 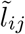 for the TF–target gene pair (*i, j*). Finally, we convert these estimated lags to interaction speeds by subtracting them from *L*, the largest possible lag handled by the trained model: 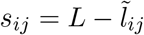

#### 5.2.6 Training details

We trained using proximal gradient descent, which minimizes the sum of the mean squared error and the hierarchical regularization. We used ReLU [62] for the activation function *σ*(·) and the identity function for the link function *s*(·). For all results, we used a single hidden layer with 16 nodes, a learning rate of 0.01, and a maximum of 10,000 epochs with early stopping as implemented by Tank et al. [58]. The hyperparameter sweep of *λ* covered 19 log-spaced values in the range [0.01, 10]. Models were implemented in PyTorch 1.12.1, and training was performed on a mix of Nvidia V100 (32 GB memory) and A100 (80 GB memory) GPUs.

#### 5.2.7 Scalability and runtime

We perform *G*Λ training runs where *G* is the number of target genes and Λ is the number of *λ* choices we sweep over. Each run is relatively light and we were able to perform 10 parallel runs for *λ* settings on a single V100 GPU. While *N* (the number of cells) is not a major runtime scalability concern, the input matrices (of size *O*(*NG*′), where *G*′ is the number of candidate regulators) need to fit into GPU memory. For large datasets (e.g., with over 100,000 cells), we recommend sketching the data [63, 64] to first extract a smaller representative sample. When the list of potential targets or regulators is known, the scope of computation can be constrained. For example, in our analysis of biological data, we limited candidate regulators to the set of transcription factors in the dataset. The (**A**′)^*l*^**X** inputs to the first hidden layer are pre-computed and amortized over all target genes, and thus this computation is not meaningful to the overall runtime. For a synthetic dataset with 1,800 cells and 100 genes, a full run of Velorama sweeping across all *λ* values required approximately 3 minutes on an Nvidia V100 GPU (**Table S7**). For scRNA-seq datasets with between 2,531– 5,780 cells, 245–499 candidate TFs, and 1,585–1,873 candidate target genes, each full run took roughly 1–3 hours on a Nvidia A100 GPU.

#### 5.2.8 Datasets and pre-processing

SERGIO We applied Velorama to four benchmark synthetic datasets obtained from SERGIO. Prior to running Velorama, we normalized the expression values of each gene to have a mean of 0 and a standard deviation of 1. To evaluate Velorama, we compared the inferred causal graph to the ground-truth set of TF–target pairs used to simulate the expression matrix. All 100 genes were considered as candidate TFs and targets.

#### 5.2.9 Single-cell RNA velocity and time series datasets

We analyzed six scRNA-seq datasets profiling mouse endocrinogenesis [30], mouse dentate gyrus neurogenesis [31], human hematopoiesis [32], cardiac hypertrophy in cardiomyocytes [27], the response of A549 cells to pracinostat [28], and human corticogenesis [42]. For the mouse endocrinogenesis, mouse dentate gyrus neurogenesis, and human hematopoiesis datasets, we used processed *Scanpy* [51] objects, which were made available by scVelo and CellRank [14, 50]. We performed a similar pre-processing procedure for each dataset. We first normalized the number of counts per cell such that each cell has a total count equal to the pre-normalization median number of counts per cell. We then applied a log(1 + *X*) transformation on the normalized counts *X*. We identified the top 2000 most highly variable genes for all datasets except for the human corticogenesis dataset, for which we identified the top 5000 most highly variable genes. The transformed expression values for each gene were then normalized to have a mean of 0 and a standard deviation of 1. We removed all genes that were represented in fewer than 1% of cells in each dataset and retained the union of the remaining highly variable genes and genes that were annotated as transcription factors according to AnimalTFDB [65]. When applying Velorama, transcription factors were selected as the set of candidate regulators, and candidate target genes were defined to be the set of highly variable genes excluding transcription factors.

#### 5.2.10 Velorama integration of time stamps and additional data modalities

Temporal information from scRNA-seq time series data can be flexibly integrated with Velorama to better inform temporal relationships between single cell measurements. scRNA-seq time series experiments typically profile populations of cells at multiple distinct time points, such that each profiled cell is associated with a coarse time stamps. We use the single-cell multimodal integration algorithm Schema [26] to endow the cells’ gene expression representations with information from these time stamps. Specifically, we use Schema to synthesize the PCA representations of cells’ expression profiles with the datasets’ time stamps, treating the former as the primary modality and the latter as the secondary modality in the Schema workflow. Schema generates transformed cell representations that unify the gene expression and time stamp modalities, and we use these new cell representations for constructing the Velorama DAG.

For multimodal single-cell data profiling both gene expression and chromatin accessibility data, we first transform the chromatin accessibility data using TF-IDF, a topic modeling approach that has been used widely in analyzing single-cell ATAC-seq data [66–68]. We apply PCA separately to each of the transform gene expression and chromatin accessibility modalities, projecting the gene expression profiles of cells onto the top *M* principal components. We exclude principle components that have Spearman correlation *ρ >* 0.9 with the total counts per cell for each of the modalities. We use Schema to synthesize the PCA representations of cells’ gene expression and chromatin accessibility profiles, designating them as the primary and secondary modalities, respectively. The resulting unified representations are then used for constructing the Velorama DAG. For both integrating time stamps and integrating chromatin accessibility data, we set the minimum correlation between the primary and secondary modalities to be 0.9 in Schema.

### 5.3 Quantification and Statistical Analysis

#### 5.3.1 TF motif accessibility analyses

The human corticogenesis dataset profiled both gene expression and chromatin accessibility in the same cell [42], presenting the opportunity to examine the relationship between the regulatory dynamics implied by its chromatin accessibility and the Velorama-inferred GRN. We applied chromVAR [43] to the chromatin accessibility data, which uses the presence (or absence) of a TF’s motif within accessible regions of the genome as a proxy for the TF’s activity per cell. We used the human position weight matrices curated by chromVAR to scan for TF motifs.

To approximate the lag between a TF’s expression and its activity, we first created pseudocells by aggregating measurements across each cell’s 100 nearest neighbors based on Euclidean distance with respect to the cells’ Schema representations. For the gene expression data, we added the raw transcript counts for each gene across each neighborhood of cells and then normalized the total counts for cell such that the total transcript counts in each cell is 10,000. For the chromatin accessibility data, we averaged the chromVAR-inferred TF activities over each cell’s neighbors. For each TF, we identify the 2% of cells with the highest expression levels and the 2% of cells with the highest chromVAR TF activity levels. We then identify the median pseudotime stamp within each of these sets of cells and compute the absolute difference between these median pseudotime associated with the TF expression and TF activity. We used this pseudotime difference as an estimate of the lag between a TF’s expression and its activity, leveraging our understanding of TF regulatory mechanisms in which changes in a TF’s expression level work together with subsequent changes in chromatin accessibility at possible genomic binding sites to drive downstream gene expression changes [69].

To evaluate the cell type-specificity of a TF’s activity, we used a *t*-test to compare the TF’s chromVAR scores within each cell type against the chromVAR scores for the same TF in all other cell types for the human corticogenesis dataset. Performing this comparison for all cell types yields a *p*-value for each cell type that represents the statistical significance of the differential TF motif accessibility for the cell type. We perform Benjamini-Hochberg correction on the *p*-values and then compute the entropy of the − log_10_(*p*) values across cell types, with lower entropy values indicating greater cell type-specificity.

#### 5.3.2 Brain-specific expression

To determine the specificity of a TF’s expression level in the human brain, we obtained consensus tissue-specific transcriptomic data from the Human Protein Atlas [70, 71]. For a TF, we calculated the mean and standard deviation of the TF’s expression levels across all non-brain tissues. Brain-specific samples were those collected from the basal ganglia, cerebellum, cerebral cortex, medulla oblongata, midbrain, pons, and white matter. We transformed the expression levels of the TF in the brain-specific tissues into a *z*-score with respect to the non-brain tisues.

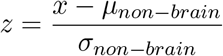

Here, *x* is the expression level for the gene in a brain-specific tissue. For each TF, we represented its brain-specific expression level as the largest *z*-score observed across the various brain-specific tissues.

## Acknowledgements

AW, RS and BB were supported by the NIH grant R35GM141861. Some figures were made using biorender.com.

## A Supplemental Information

### A.1 Velorama Architecture

**Supplementary Figure S1:**
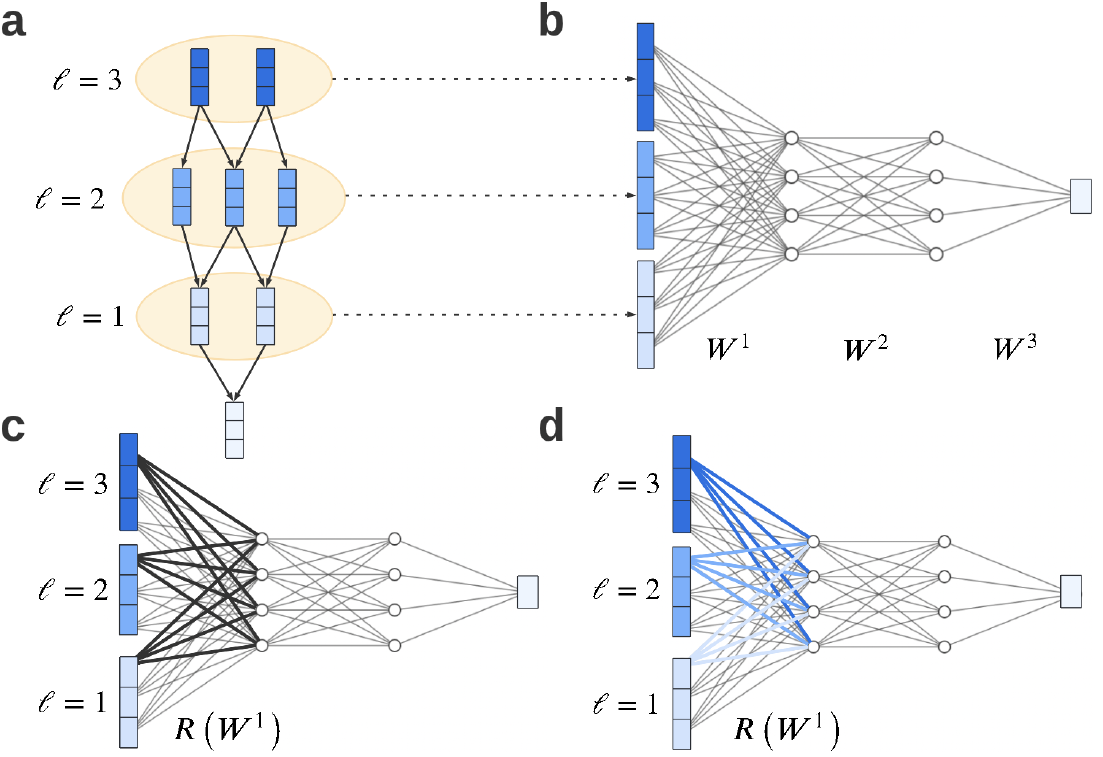
Velorama architecture: *N* = 8, *G* = 3, *L* = 3, *d* = 4 and *K* = 3. (a) Velorama predicts a node’s features based on ancestors in the DAG. (b) Predictions are made by passing aggregated information from each lag through a feed-forward neural network. (c) If the weights associated with variable *i* in *fj* (highlighted in black for *i* = 1) are all equal to 0, then variable *i* does *not* Granger-cause variable *j*. (d) The hierarchical penalty further penalizes groups of weights associated with particular lags.

### A.2 Velorama-inferred GRNs for SERGIO datasets

We plotted the ground-truth GRN as well as the Velorama-inferred GRNs using pseudotime and RNA velocity for datasets A, B and C (**Figure S2**). Using RNA velocity improves predictions of the GRN.

**Supplementary Figure S2:**
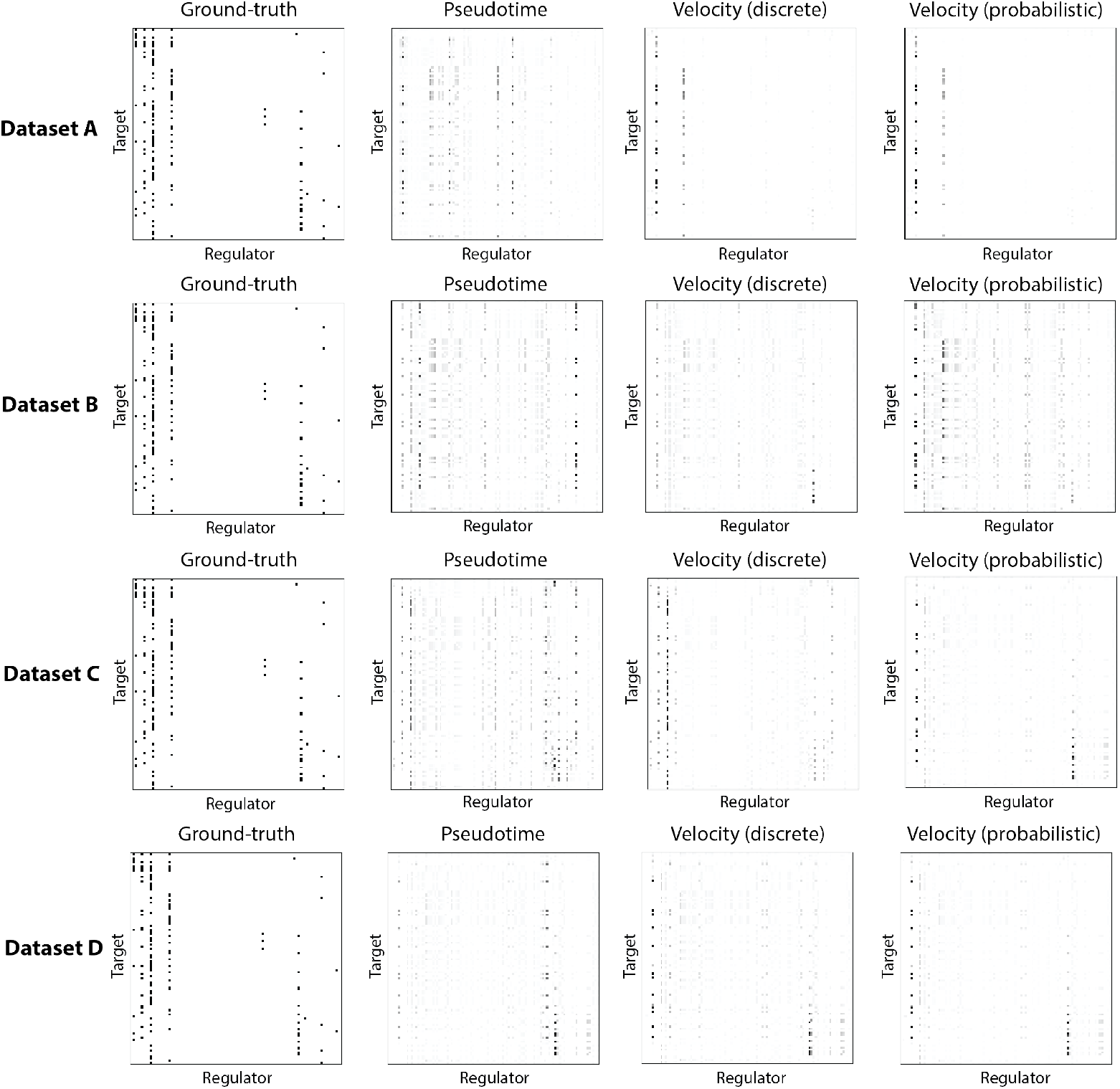
Ground-truth GRN for Datasets **A** (top), **B** (middle) and **C** (bottom) compared to the Velorama-inferred GRNs when using pseudotime or RNA velocity data.

### A.3 Velorama time stamp integration and GRN inference accuracy on SERGIO datasets

SERGIO’s default settings do not produce time stamps for each cell, but we can manually assign time stamps that cohere with SERGIO’s procedure for simulating differentiation trajectories. For a given differentiation trajectory, a cell type transition is initiated by changing the production rates of the underlying GRN’s master regulators, after which SERGIO’s time-course simulations are applied to generate the downstream expression changes that follow from these modified rates. As a result, the cell type transition points demarcate the boundaries of the distinct time windows that are used in the simulation, and each cell type’s expression profiles directly corresponds to one of these time windows. For a linear trajectory like Dataset A, each cell type would be assigned a distinct time stamp (*t* = 0, 1, 2) (**Figure 3A**). For branching trajectories, the cell types that are differentiated from the same parent cell type exist at the same time point, and we mark each cell from a given cell type with the same time stamp. For example, for Dataset C, the time stamps assigned to each cell type would be as follows: red (*t* = 0), green (*t* = 1), blue (*t* = 2), orange (*t* = 3), purple (*t* = 3), yellow (*t* = 3) (**Figure 3A**).

**Table S1:** Velorama performance on SERGIO datasets with and without integrating time stamps.

| Method | Dataset A |  | Dataset B |  | Dataset C |  | Dataset D |  |
| --- | --- | --- | --- | --- | --- | --- | --- | --- |
|  | AUPRC | AUROC | AUPRC | AUROC | AUPRC | AUROC | AUPRC | AUROC |
| Velorama (pseudotime) | 0.191 | 0.848 | 0.154 | 0.856 | 0.327 | 0.882 | 0.296 | 0.872 |
| Velorama (pseudotime + time stamps) | 0.112 | 0.894 | 0.482 | 0.912 | 0.592 | 0.951 | 0.672 | 0.939 |
| Velorama (velocity) | 0.399 | 0.900 | 0.516 | 0.856 | 0.596 | 0.923 | 0.541 | 0.926 |
| Velorama (velocity + time stamps) | 0.298 | 0.901 | 0.717 | 0.952 | 0.691 | 0.963 | 0.738 | 0.958 |

### A.4 Velorama prioritizes temporally causal regulatory relationships

To demonstrate that Velorama prioritizes TF–target interactions that are temporally causal, we examined Velorama’s predictions for the mouse dentate gyrus neurogenesis dataset [31]. We focused on this dataset because it features true time stamps that can be used to identify TF–target interactions that align with the temporal flow of causal regulation, in which activation (i.e. upregulation) of a TF occurs at an earlier time stamp than the change in expression of its downstream target gene. In particular, we evaluated the enrichment of such TF–target interactions in Velorama’s predictions. We found that these temporally causal TF–target gene interactions were indeed more strongly enriched in Velorama’s predictions than interactions that do not display this temporally causal relationship.

**Table S2:**
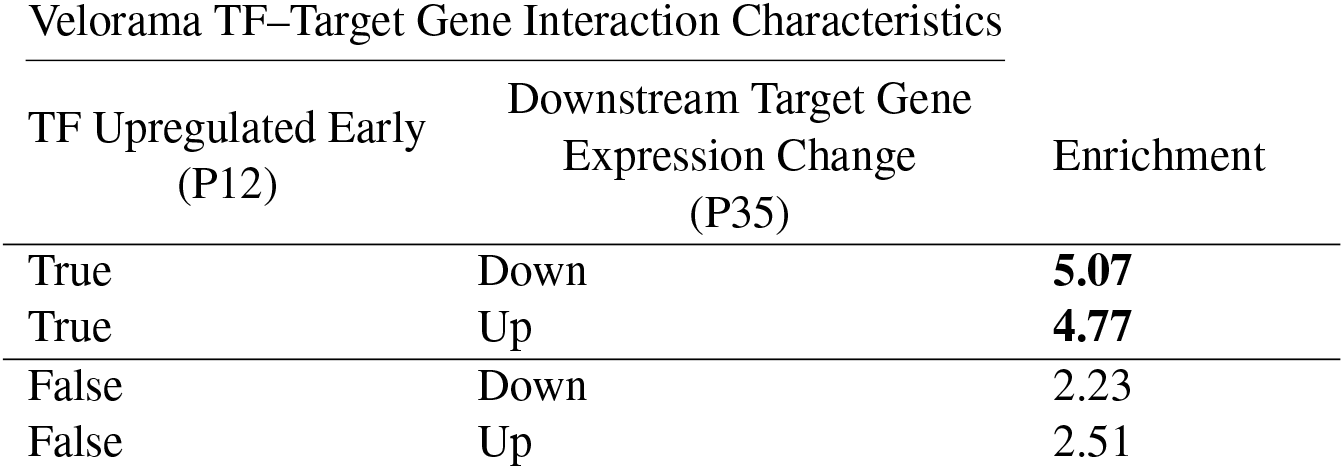
Enrichment of various categories of TF–target gene interactions inferred by Velorama.

| Velorama TF–Target Gene Interaction Characteristics |  |  |
| --- | --- | --- |
| TF Upregulated Early<br>(P12) | Downstream Target Gene<br>Expression Change<br>(P35) | Enrichment |
| True | Down | **5.07** |
| True | Up | **4.77** |
| False | Down | 2.23 |
| False | Up | 2.51 |

### A.5 Overlap of TF–target gene interactions inferred using Velorama and scDiff

scDiff [25] leverages time stamps from single-cell time series data to infer driver TFs and their downstream target genes at each time point. Using the mouse dentate gyrus neurogenesis dataset [31], which profiled cells at multiple time points, we compared the TF–target gene interactions inferred using Velorama and RNA velocity with those of scDiff, which explicitly uses the data’s time stamps. To directly compare Velorama’s predictions and those of scDiff, we compared the TF–target gene interactions featuring TFs and targets retained in the GRNs inferred by both methods.

**Supplementary Figure S3:**
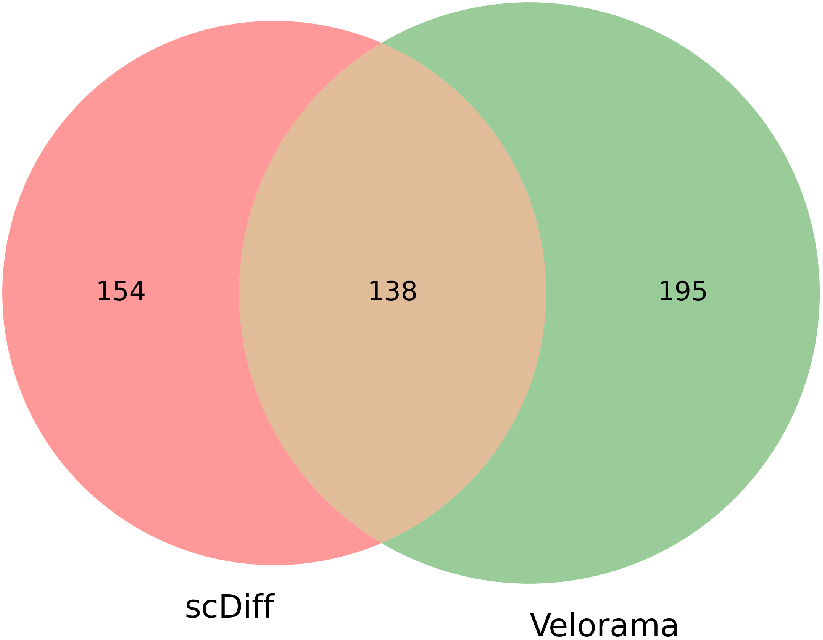
Venn diagram of TF–target gene interactions inferred by Velorama and scDiff

### A.6 RNA velocity coheres with true time stamps

To evaluate the coherence of RNA velocity with true time stamps from single-cell time series data, we analyzed the CellRank cell–cell transition probabilites inferred for the mouse dentate gyrus neurogenesis dataset [31], which profiled cells at multiple time points. We focused on the cell–cell transitions that feature cells from distinct time points and calculated the estimated probabilities for the set of cell–cell transitions that align with the time stamps (e.g., transition from P12 to P35) and those that misalign with the time stamps (e.g., transition from P35 to P12).

**Supplementary Figure S4:**
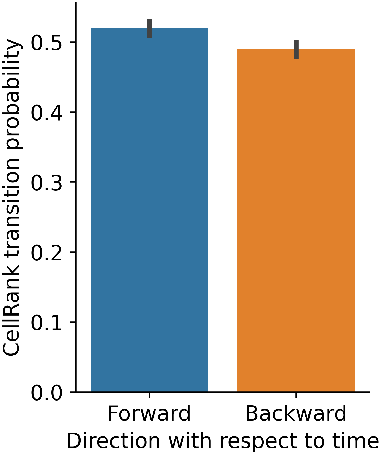
CellRank probabilities for cell–cell transitions that align vs. misalign with true time stamps

### A.7 Velorama-inferred TF speed estimates are anti-correlated with TF expression-activity lags

**Supplementary Figure S5:**
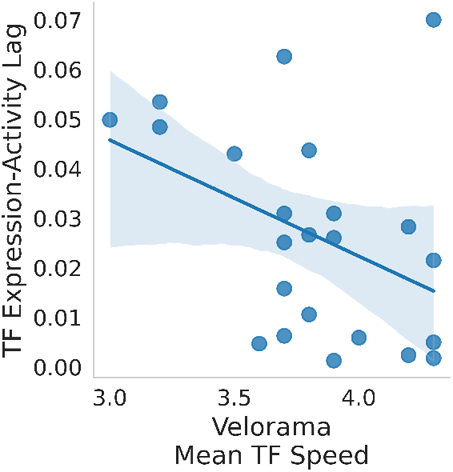
Relationship between Velorama-inferred TF speed estimates and TF expression-activity lags (Pearson’s *ρ* = −0.41, *p* = 0.05).

### A.8 Velorama robustness to hyperparameter choices

**Table S3:**
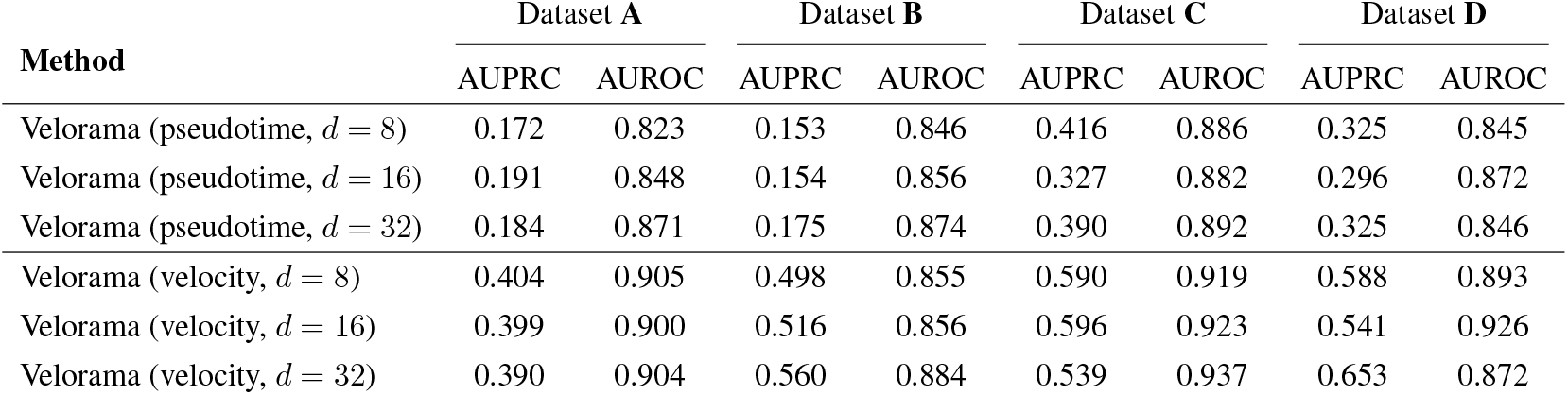
Velorama performance on SERGIO datasets across different choices of number of hidden dimensions *d*.

| Method | Dataset A |  | Dataset B |  | Dataset C |  | Dataset D |  |
| --- | --- | --- | --- | --- | --- | --- | --- | --- |
|  | AUPRC | AUROC | AUPRC | AUROC | AUPRC | AUROC | AUPRC | AUROC |
| Velorama (pseudotime, $d = 8$) | 0.172 | 0.823 | 0.153 | 0.846 | 0.416 | 0.886 | 0.325 | 0.845 |
| Velorama (pseudotime, $d = 16$) | 0.191 | 0.848 | 0.154 | 0.856 | 0.327 | 0.882 | 0.296 | 0.872 |
| Velorama (pseudotime, $d = 32$) | 0.184 | 0.871 | 0.175 | 0.874 | 0.390 | 0.892 | 0.325 | 0.846 |
| Velorama (velocity, $d = 8$) | 0.404 | 0.905 | 0.498 | 0.855 | 0.590 | 0.919 | 0.588 | 0.893 |
| Velorama (velocity, $d = 16$) | 0.399 | 0.900 | 0.516 | 0.856 | 0.596 | 0.923 | 0.541 | 0.926 |
| Velorama (velocity, $d = 32$) | 0.390 | 0.904 | 0.560 | 0.884 | 0.539 | 0.937 | 0.653 | 0.872 |

**Table S4:** Velorama performance on SERGIO datasets across different choices of number of nearest neighbors *k*.

| Method | Dataset A |  | Dataset B |  | Dataset C |  | Dataset D |  |
| --- | --- | --- | --- | --- | --- | --- | --- | --- |
|  | AUPRC | AUROC | AUPRC | AUROC | AUPRC | AUROC | AUPRC | AUROC |
| Velorama (pseudotime, $k = 15$) | 0.191 | 0.848 | 0.154 | 0.856 | 0.327 | 0.882 | 0.296 | 0.872 |
| Velorama (pseudotime, $k = 30$) | 0.191 | 0.848 | 0.168 | 0.862 | 0.384 | 0.885 | 0.323 | 0.845 |
| Velorama (velocity, $k = 15$) | 0.399 | 0.900 | 0.516 | 0.856 | 0.596 | 0.923 | 0.541 | 0.926 |
| Velorama (velocity, $k = 30$) | 0.399 | 0.900 | 0.556 | 0.880 | 0.489 | 0.942 | 0.552 | 0.925 |

**Table S5:** Velorama performance on SERGIO datasets across different choices of number of PCs *M*.

| Method | Dataset A |  | Dataset B |  | Dataset C |  | Dataset D |  |
| --- | --- | --- | --- | --- | --- | --- | --- | --- |
|  | AUPRC | AUROC | AUPRC | AUROC | AUPRC | AUROC | AUPRC | AUROC |
| Velorama (pseudotime, $M = 30$) | 0.127 | 0.875 | 0.137 | 0.852 | 0.161 | 0.893 | 0.394 | 0.879 |
| Velorama (pseudotime, $M = 50$) | 0.191 | 0.848 | 0.168 | 0.862 | 0.384 | 0.885 | 0.323 | 0.845 |
| Velorama (velocity, $M = 30$) | 0.381 | 0.888 | 0.521 | 0.866 | 0.484 | 0.932 | 0.501 | 0.940 |
| Velorama (velocity, $M = 50$) | 0.399 | 0.900 | 0.556 | 0.880 | 0.489 | 0.942 | 0.552 | 0.925 |

**Table S6:**
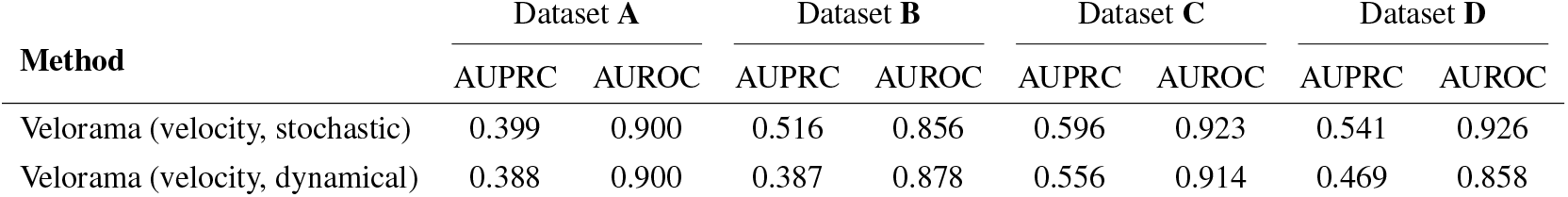
Velorama performance on SERGIO datasets across different choices of RNA velocity dynamical models.

| Method | Dataset A |  | Dataset B |  | Dataset C |  | Dataset D |  |
| --- | --- | --- | --- | --- | --- | --- | --- | --- |
|  | AUPRC | AUROC | AUPRC | AUROC | AUPRC | AUROC | AUPRC | AUROC |
| Velorama (velocity, stochastic) | 0.399 | 0.900 | 0.516 | 0.856 | 0.596 | 0.923 | 0.541 | 0.926 |
| Velorama (velocity, dynamical) | 0.388 | 0.900 | 0.387 | 0.878 | 0.556 | 0.914 | 0.469 | 0.858 |

### A.9 Runtimes of GRN inference algorithms on SERGIO datasets

**Table S7:** GRN inference runtimes (in minutes)

| Method | Dataset A | Dataset B | Dataset C | Dataset D |
| --- | --- | --- | --- | --- |
| SINGE | 49.9 | 87.5 | 183.9 | 248.7 |
| SINCERITIES | 0.2 | 0.2 | 0.2 | 0.2 |
| SCODE | 9.1 | 10.4 | 13.2 | 14.5 |
| PIDC | 0.3 | 0.3 | 0.4 | 0.4 |
| Scribe | 3.9 | 5.3 | 9.2 | 12.9 |
| GRNBoost2 | 3.3 | 5.4 | 11.5 | 10.3 |
| GENIE3 | 6.9 | 9.2 | 14.1 | 17.3 |
| RNA-ODE | 12.9 | 17.3 | 25.5 | 32.5 |
| Velorama | 2.5 | 2.6 | 3.3 | 2.9 |

